# Total-evidence phylogenetic analysis resolves the evolutionary timescale of mantis shrimps (Stomatopoda) and provides insights into their molecular and morphological evolutionary rates

**DOI:** 10.1101/2023.11.05.565425

**Authors:** Cara Van Der Wal, Shane T. Ahyong, Maxim W.D. Adams, Nathan Lo, Simon Y.W. Ho

**Affiliations:** School of Life and Environmental Sciences, University of Sydney, Sydney, NSW 2006, Australia; Australian Museum Research Institute, Australian Museum, 1 William Street, Sydney, NSW 2000, Australia; School of Biological, Earth and Environmental Sciences, University of New South Wales, Kensington, NSW 2052, Australia

**Keywords:** crustacean, Bayesian phylogenetics, molecular clock, evolutionary rate, morphology

## Abstract

The crustacean order Stomatopoda comprises approximately 500 species of mantis shrimps. These marine predators, common in tropical and subtropical waters, possess sophisticated visual systems and specialized hunting appendages. In this study, we infer the evolutionary relationships within Stomatopoda using a combined data set of 77 morphological characters, whole mitochondrial genomes, and three nuclear markers. Our data set includes representatives from all seven stomatopod superfamilies, including the first sequence data from Erythrosquilloidea. Using a Bayesian relaxed molecular clock with fossil-based calibration priors, we estimate that crown-group unipeltatan stomatopods appeared ∼140 (95% credible interval 201–102) million years ago in the Mesozoic. Additionally, our results support the hypothesis that specialized smashing and spearing appendages appeared early in the evolutionary history of Unipeltata. We found no evidence of a correlation between rates of morphological and molecular evolution across the phylogeny, but identified very high levels of among-lineage rate variation in the morphological characters. Our total-evidence analysis recovered evolutionary signals from both molecular and morphological data sets, demonstrating the merit in combining these sources of information for phylogenetic inference and evolutionary analysis.

## 1. Introduction

Stomatopod crustaceans (Malacostraca: Stomatopoda), or mantis shrimps, are known for their advanced visual systems and their unique ‘spearing’ and ‘smashing’ hunting appendages (Marshall et al., 2007; Patek, 2019). Previous studies of stomatopods have focused on these novel features, as well as their charismatic nature and social behaviors (Gagnon et al., 2015; Green and Patek, 2018; Franklin et al., 2019; Koga and Rouse, 2021). Despite this research, the phylogenetic relationships among the major lineages of stomatopods have not been resolved with confidence. The approximately 500 extant species of stomatopods are divided into seven superfamilies: Gonodactyloidea, Squilloidea, Lysiosquilloidea, Bathysquilloidea, Erythrosquilloidea, Parasquilloidea, and Eurysquilloidea (Ahyong, 1997, 2001). The three largest and best studied superfamilies are Gonodactyloidea, Squilloidea, and Lysiosquilloidea (Ahyong and Harling, 2000; Ahyong and Jarman, 2009). The extant stomatopods form the suborder Unipeltata, while the order also includes the extinct suborders Palaeostomatopodea and Archaeostomatopodea (Schram et al., 2013).

Studies of stomatopod morphology provide the basis for the current classification of the group (Manning, 1968, 1980; Ahyong, 1997; Ahyong and Harling, 2000). Analyses of morphological data provide strong support for some of the relationships among superfamilies, including the placements of Squilloidea and Lysiosquilloidea; however, they have been unable to robustly resolve other deep nodes in the tree, such as the position of Bathysquilloidea (Ahyong and Harling, 2000). Recent molecular studies have shed light on the evolutionary relationships and divergence times among the stomatopod superfamilies, but have produced phylogenetic estimates with generally weak support for deep nodes. Porter et al. (2010) and Van Der Wal et al. (2017) analysed four and five loci, respectively, to infer a sister relationship between Gonodactyloidea and Squilloidea. In contrast, Ahyong and Jarman (2009) found a sister relationship between Lysiosquilloidea and Squilloidea in their analysis of three loci. The poor phylogenetic resolution of deep stomatopod relationships has persisted in analyses of whole mitochondrial genomes (Hwang and Jung, 2021; Hwang et al., 2021; Koga and Rouse, 2021; Yang et al., 2021), while the placement of the superfamily Erythrosquilloidea has not been interrogated using molecular data.

Combined molecular and morphological data sets provide valuable opportunities for improving phylogenetic support, as well as understanding the broad relationship between evolutionary processes at the genetic and phenotypic scales. For example, a correlation between morphological and molecular evolutionary rates might be expected if we assume that changes in phenotype are generally accompanied by genetic change (Omland, 1997; Seligmann, 2010; Asar et al., 2023). An analysis of a large data set detected a correlation between rates of morphological and molecular evolution in crustaceans and other arthropods during the Cambrian explosion (Lee et al., 2013). Ecological opportunism or smaller body size, coupled with short generation times, was proposed to explain this correlation. However, there have been few studies of molecular and morphological evolutionary rates in stomatopods. Studies of stomatopods found no evidence that sea surface temperature, depth, latitude, and habitat type influence the rate of morphological evolution (Reaka and Manning, 1981, 1987). However, they did find evidence for an inverse relationship between body size and rates of morphological evolution. In contrast, body size does not show a clear association with rates of molecular evolution in invertebrates (Thomas et al., 2006).

In this study, we use a combination of molecular and morphological data to infer an evolutionary time-tree for extant stomatopods. Our total-evidence analyses are based on 77 morphological characters, whole mitochondrial genomes, and three nuclear genes from representatives of all seven superfamilies. We present a new timescale of evolution for the group, including a comparison of morphological and molecular evolutionary rates. Our results demonstrate the evolutionary insights that can be gained by using combined data sets for phylogenetic inference.

## 2. Materials and methods

### 2.1. Taxon sampling

Tissue samples were collected from 26 stomatopod species preserved in ethanol or formalin from the Australian Museum (Sydney), Florida Museum of Natural History (Gainesville), Lee Kong Chian Natural History Museum (National University of Singapore), Muséum national d’Histoire naturelle (Paris), and Western Australian Museum (Perth). Genetic sequence data from these samples were combined with publicly available data on GenBank for a focal set of 34 taxa. Our data set includes representatives of all seven stomatopod superfamilies (Bathysquilloidea *n* = 3, Erythrosquilloidea *n* = 1, Eurysquilloidea *n* = 2, Gonodactyloidea *n* = 10, Lysiosquilloidea *n* = 7, Parasquilloidea *n* = 1, and Squilloidea *n* = 10). These taxa represent 30 of the 118 stomatopod genera, belonging to 16 of the 17 stomatopod families (Supplementary Tables S1 and S2).

### 2.2. Morphological characters

We scored the morphology of the 34 stomatopod exemplars in our data set (Supplementary Tables S3 and S4). Our morphological matrix includes 77 unordered and equally weighted variable characters described by Ahyong and Harling (2000). These characters were found to be phylogenetically informative by Ahyong and Harling (2000) and are derived from external somatic morphology, in particular, features of the carapace, abdomen, telson, eyes, maxillipeds, and pleopods.

The morphological character matrix was analysed using both maximum likelihood and Bayesian inference. Maximum-likelihood phylogenetic estimates were obtained using RAxML 8.0.14 (Stamatakis, 2014). To account for the lack of invariant sites in the morphological data, we used the Lewis-type correction for ascertainment bias (Lewis, 2001). Bootstrapping was used to quantify nodal support for the estimated topology, based on 1000 pseudoreplicates of the data.

Bayesian phylogenetic analysis was performed using MrBayes 3.2.5 (Ronquist et al., 2012), with the Mkv model of character change. The posterior distribution was estimated using Markov chain Monte Carlo (MCMC) sampling, with one cold and three heated Markov chains. We drew samples every 10^3^ steps over a total of 10^7^ MCMC steps and discarded the initial 25% of samples as burn-in. The analysis was run in duplicate and convergence was evaluated using the average standard deviation of split frequencies. Samples from the two runs were combined and the maximum-clade-credibility tree was identified using TreeAnnotator, part of the BEAST 2 package (Bouckaert et al., 2019).

### 2.3. Mitochondrial and nuclear data

We obtained gill and muscle tissue samples from 26 stomatopods for DNA extraction. Genomic DNA was extracted from 22 ethanol-preserved specimens using the Qiagen DNeasy Blood & Tissue Kit (Qiagen, Hilden, Germany), and from four formalin-fixed specimens using the QIAamp DNA FFPE Tissue Kit (Qiagen, Hilden, Germany) following the manufacturer’s instructions. The genomic DNA samples were quantified using a Qubit 2.0 Fluorometer (Thermo Fisher Scientific, Sydney, Australia) and sent to Macrogen (Seoul, South Korea) and BGI (Shenzhen, China) for shotgun sequencing. Paired-end sequencing was performed at both facilities (150 bp), with Macrogen and BGI using the Illumina X-Ten and Illumina HiSeq 2500 platforms, respectively. Using these sequencing platforms, we generated 1–2 Gb of shotgun reads.

To assemble the mitochondrial genomes, we generated BLAST databases to filter the reads according to their similarity to other crustacean mitochondrial genomes. We performed *de novo* assemblies on the filtered set of reads for each species using the Qiagen CLC Genomics Workbench 10 (https://www.qiagenbioinformatics.com). The assembled contigs for each species were then mapped to a reference genome (≥4× coverage) to obtain consensus sequences in Geneious 10.2.3 (https://www.geneious.com). To map contigs we used the published mitochondrial reference genome of *Oratosquilla oratoria* (de Haan, 1844) (Liu and Cui, 2010). The 26 mitochondrial genomes were annotated using the online MITOS WebServer (Bernt et al., 2013). Most of the mitochondrial protein-coding genes were recovered in all our samples, but some of the highly degraded specimens (formalin fixed) yielded incomplete sequences. For each species, we concatenated the protein-coding genes, transfer RNA genes (tRNAs), and ribosomal RNA genes (rRNAs).

A similar approach was taken to assemble nuclear sequence data. We generated a BLAST database to identify and filter reads from three nuclear genes: small subunit 18S rRNA (*18S*), large subunit 28S rRNA (*28S*), and histone 3 (*H3*). We selected these three genes because of their online availability and effectiveness in resolving phylogenetic relationships. We performed *de novo* assemblies on the filtered set of reads for each species and each nuclear gene using the CLC Genomics Workbench 10 and mapped the contigs to reference sequences, as described above for the mitochondrial genomes.

The sequences of the mitochondrial protein-coding genes, mitochondrial tRNAs, mitochondrial rRNAs, and nuclear markers were aligned separately using MUSCLE 3.8.31 (Edgar, 2004). The nucleotide sequences of the protein-coding genes were translated to amino acids to check for frameshift mutations and premature stop codons. Given the potentially negative impacts of substitution saturation on phylogenetic inference, we applied Xia’s test in DAMBE 6 (Xia, 2017) to the tRNA genes, each rRNA gene, and each nuclear gene, and each of the three codon sites of the mitochondrial protein-coding genes. We found evidence of saturation in mitochondrial *16S* and third codon sites, based on simulations on a symmetrical tree (Supplementary Table S5). Therefore, we removed these subsets of the data for all subsequent phylogenetic analyses. The remaining alignments were concatenated for further analyses (total length of 14,598 bp). We found that the degree of compositional heterogeneity in the data set posed a low risk of causing biased phylogenetic inferences, as evaluated using PhyloMAd (Duchêne et al., 2017, 2018). The optimal partitioning scheme was selected for the data set using the Bayesian information criterion in PartitionFinder 2.1.1 (Lanfear et al., 2012; Lanfear et al., 2017), and implemented in all of our phylogenetic analyses (Supplementary Table S6).

We analysed the concatenated data set using maximum likelihood and Bayesian inference. Maximum-likelihood analysis was performed using RAxML with 1000 bootstrap replicates, with a separate GTR+G model of nucleotide substitution applied to each data subset. Bayesian phylogenetic estimates were obtained using MrBayes, with the same partitioning scheme and substitution models as used in our maximum-likelihood analysis. MCMC sampling was performed with one cold and three heated Markov chains, with samples drawn every 2×10^3^ steps over a total of 2×10^7^ MCMC steps. The initial 25% of samples were discarded as burn-in. The analysis was run in duplicate and convergence was evaluated using the average standard deviation of split frequencies.

### 2.4. Combined morphological and molecular data

We assembled a concatenated data set containing 77 morphological characters and 14,598 nucleotide sites. Total-evidence phylogenetic analyses were performed using both maximum likelihood and Bayesian inference. The partitioning scheme for the molecular data followed the recommendation of PartitionFinder as described above, but with an additional subset for the morphological data (Supplementary Table S6). Maximum-likelihood analyses were performed using RAxML with 1000 bootstrap replicates. The Mkv model of character change was applied to the morphological characters, whereas a separate GTR+G model was assigned to each of the eight molecular data subsets.

To co-estimate the stomatopod phylogeny and evolutionary timescale, we conducted Bayesian phylogenetic analyses in BEAST 2.6.4 (Bouckaert et al., 2019) using the same data-partitioning scheme as in our maximum-likelihood analyses. We used Bayesian model averaging (bModelTest) for the substitution model (Bouckaert and Drummond, 2017), allowing a separate evolutionary rate for each of the data subsets (Duchêne et al., 2020). To investigate the relationship between rates of morphological and molecular evolution, we ran two analyses: the first using separate uncorrelated log-normal clock models for the morphological and molecular data sets, and the second using separate uncorrelated log-normal clock models for the morphological, mitochondrial, and nuclear data sets (Drummond et al., 2006). Owing to the composition of our morphological data set, and because we were primarily interested in the evolutionary rates within Stomatopoda, we did not include outgroup taxa. Instead, we placed a monophyly constraint on all taxa excluding *Hemisquilla*, based on the highly supported position of *Hemisquilla* as the sister lineage of all other extant stomatopods in most previous phylogenetic analyses of the group (Ahyong and Jarman, 2009; Porter et al., 2010; Van Der Wal et al., 2017). A recent analysis of mitochondrial genomes found a nested position of *Hemisquilla* within Stomatopoda (Koga and Rouse, 2021); topology tests showed some support for this placement, but the inclusion of highly genetically divergent outgroup taxa might have had a disruptive impact on phylogenetic inference.

The molecular dating analyses were calibrated using fossil evidence. Each fossil calibration was implemented as an exponential prior on the corresponding node age (Supplementary Table S7), reflecting a declining probability of the node age being older than the minimum age indicated by the fossil evidence (Nguyen and Ho, 2020). The 97.5% soft maximum for the age of crown stomatopods was based on the appearance of *Daidal acanthocercus* Jenner, Hof & Schram, 1998 (Stomatopoda, Archaeostomatopodea) in the Carboniferous (313 million years ago, Ma) (Schram, 2007; Van Der Wal et al., 2017). The occurrence ages of five stomatopod fossils were used to calibrate internal nodes. We used fossils reliably assigned to four extant genera (*Pseudosquilla*, *Odontodactylus*, *Bathysquilla*, and *Squilla*) and one extinct stomatopod genus (Squilloidea: *Ursquilla*) (Haug et al., 2013; Schram et al., 2013; Wolfe et al., 2016; Van Der Wal et al., 2017). The four fossils corresponding to extant genera were used to calibrate the split between each genus and its sister lineage. The *Ursquilla* fossil was used to calibrate the node separating Squilloidea and *Alainosquilla* (Gonodactyloidea) (Supplementary Table S7).

To evaluate the sensitivity of the results to the choice of tree prior, we conducted analyses using both a Yule speciation model and a birth-death model for the tree prior (Stadler, 2009). These two models were compared using marginal likelihoods, calculated with the stepping-stone estimator (Xie et al., 2011), and the Bayes factor was interpreted according to the guidelines of Kass and Raftery (1995). Posterior distributions of parameters were estimated using MCMC sampling, with samples drawn every 10^4^ steps over a total of 10^8^ steps. We ran analyses in duplicate to check for MCMC convergence and inspected the samples using Tracer 1.7.1 (Rambaut et al., 2018). To compare morphological and molecular evolutionary rates, we tested for a correlation using the Bayesian posterior mean estimates of branch rates (Asar et al., 2023).

## 3. Results

### 3.1. Separate analyses of morphological and molecular data

The separate phylogenetic analyses of the morphological and molecular data sets produced substantially different estimates of the stomatopod phylogeny. Maximum-likelihood and Bayesian analyses of the morphological data yielded estimates that had generally low node support (Fig. 1). When rooted along the branch leading to Hemisquillidae (*Hemisquilla*), the morphological phylogeny placed the clade comprising Squilloidea, Parasquilloidea, Eurysquilloidea, and Bathysquilloidea as a well-supported sister group to Lysiosquilloidea and Erythrosquilloidea. Lysiosquilloidea was not found to be monophyletic, owing to the nested position of Erythrosquilloidea. Our results support the monophyletic origin of specialized smashing in stomatopods (all gonodactyloids).

**Fig. 1.**
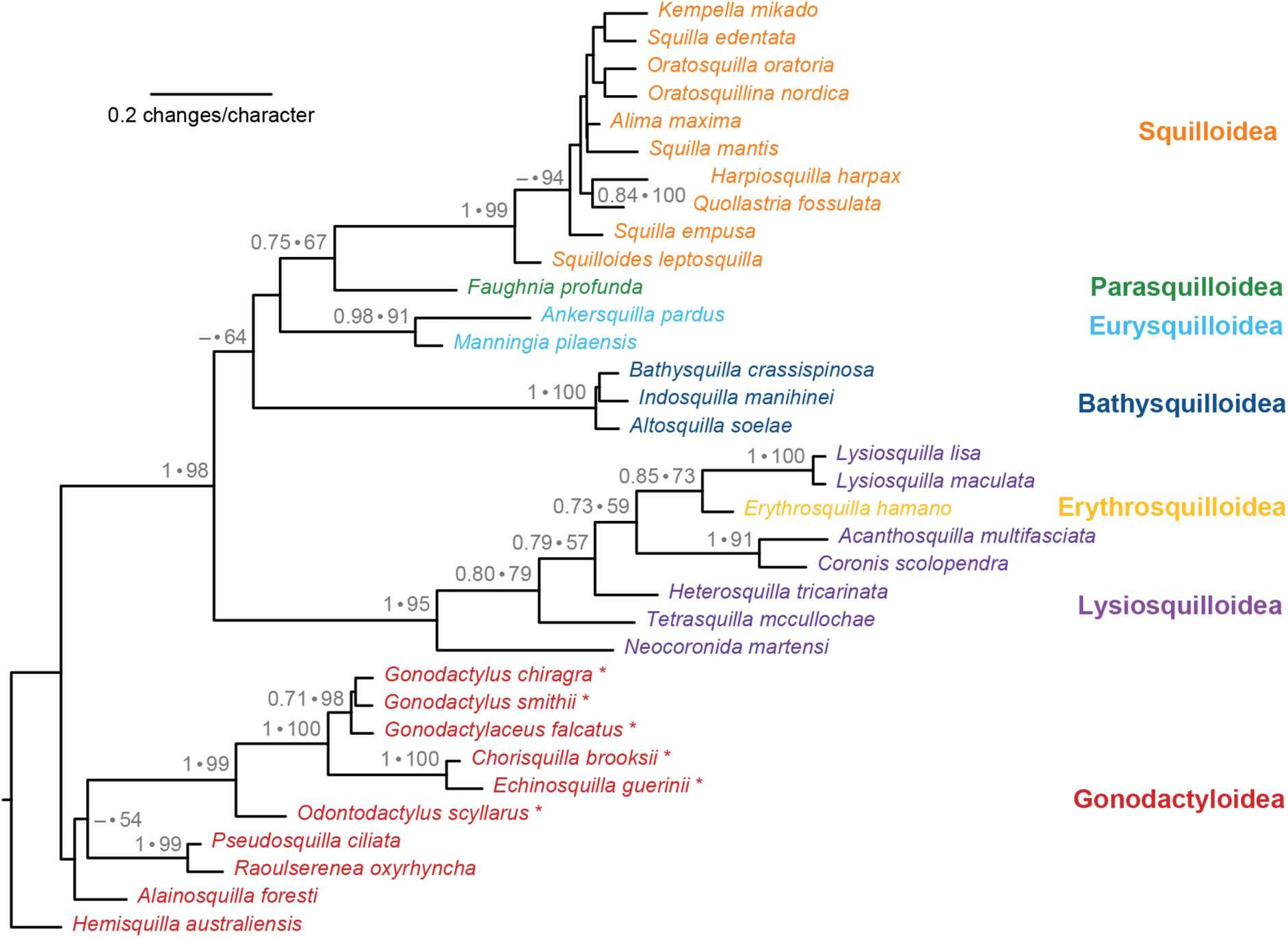
Bayesian maximum-clade-credibility phylogeny of the seven stomatopod superfamilies inferred using 77 morphological characters. Support values at nodes are shown for posterior probabilities above 0.50 and likelihood bootstrap percentages above 50%. Taxon colors correspond to their superfamily classification. Asterisks indicate species with specialized ‘smashing’ appendages.

In contrast, the tree inferred from molecular data, when rooted on the branch leading to Hemisquillidae (*Hemisquilla*), placed the gonodactyloid family Pseudosquillidae as the sister clade to the remaining stomatopods with moderately high support (Fig. 2). The inferred relationships among the three major superfamilies also differed from those estimated from morphological data. In a broad sense, the molecular data supported Squilloidea as the sister group to a clade containing Lysiosquilloidea, Gonodactyloidea (excluding Hemisquillidae and Pseudosquillidae), Erythrosquilloidea, and Bathysquilloidea. In agreement with the tree inferred from morphological data, Erythrosquilloidea was nested within a paraphyletic Lysiosquilloidea. A sister relationship between Bathysquilloidea and a clade comprising Lysiosquilloidea and Erythrosquilloidea was strongly supported. However, there was weak support for deep nodes in the molecular phylogeny, rendering it difficult to draw conclusions about the relationships among superfamilies. Smashing stomatopods, comprising the majority of gonodactyloids, formed a monophyletic group that was nested within a paraphyletic group of spearing stomatopods (Fig. 2).

**Fig. 2.**
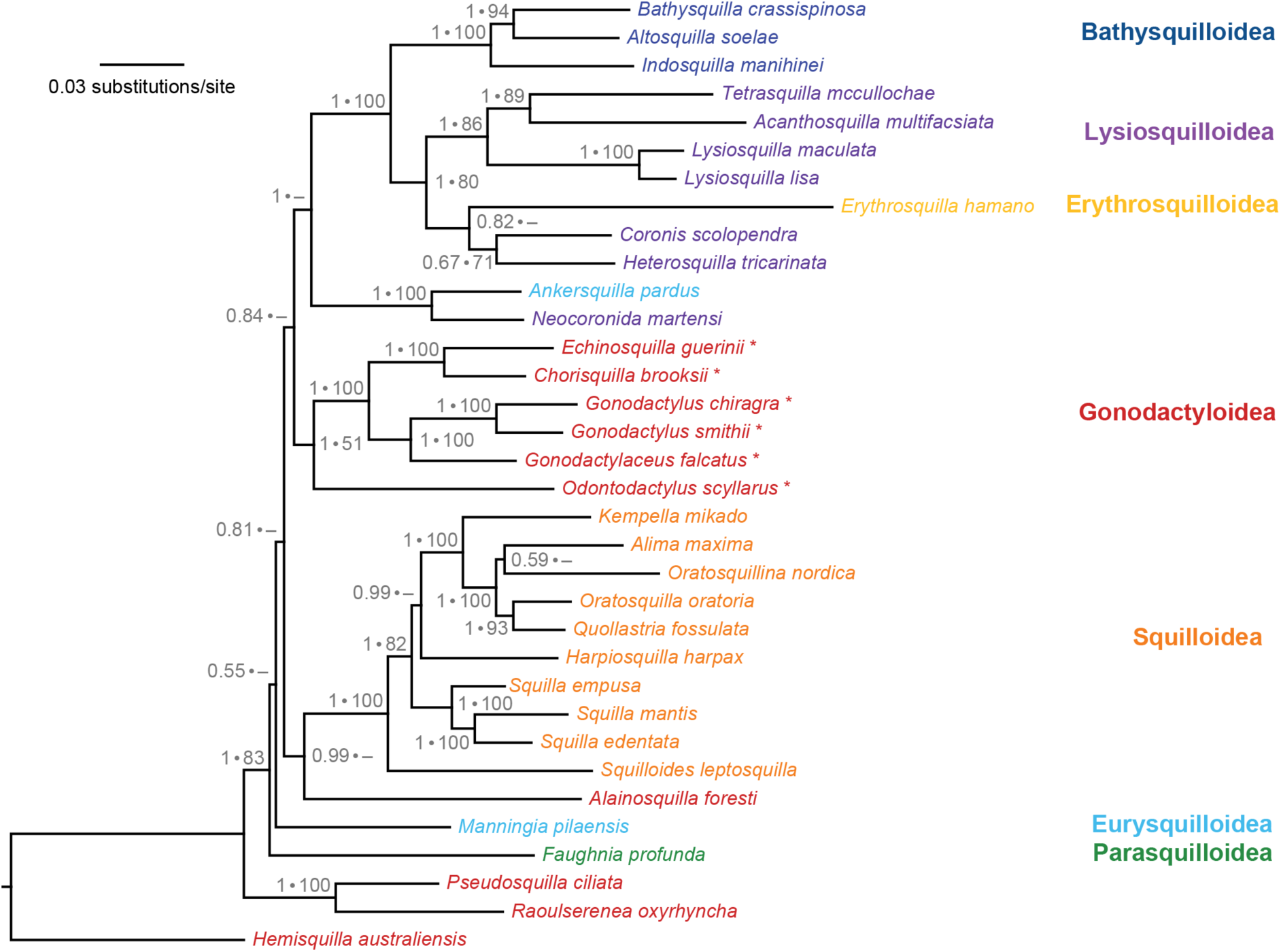
Bayesian maximum-clade-credibility phylogeny of the seven stomatopod superfamilies inferred from mitochondrial genomes and nuclear *18S*, *28S*, and *H3* (14,598 bp). Support values at nodes are shown for posterior probabilities above 0.50 and likelihood bootstrap percentages above 50%. Taxon colors correspond to their superfamily classification. Asterisks indicate species with specialized ‘smashing’ appendages.

### 3.2. Total-evidence analyses and evolutionary rates

Maximum-likelihood and Bayesian analyses yielded almost identical phylogenetic estimates for the combined morphological and molecular data set. The total-evidence phylogenetic estimate was generally well supported, including the deep nodes of the tree (Fig. 3). The tree supported Pseudosquillidae as the sister group to the remaining stomatopods (excluding *Hemisquilla*), which is consistent with the results from our analysis of the molecular data alone. The relationships among the three major superfamilies resemble those in the tree inferred from morphological data alone: Gonodactyloidea (excluding *Hemisquilla*, Pseudosquillidae, and *Alainosquilla*) forms the sister clade to the remaining superfamilies. Three superfamilies, Lysiosquilloidea, Gonodactyloidea, and Eurysquilloidea, as currently defined, were not found to be monophyletic (Fig. 3).

**Fig. 3.**
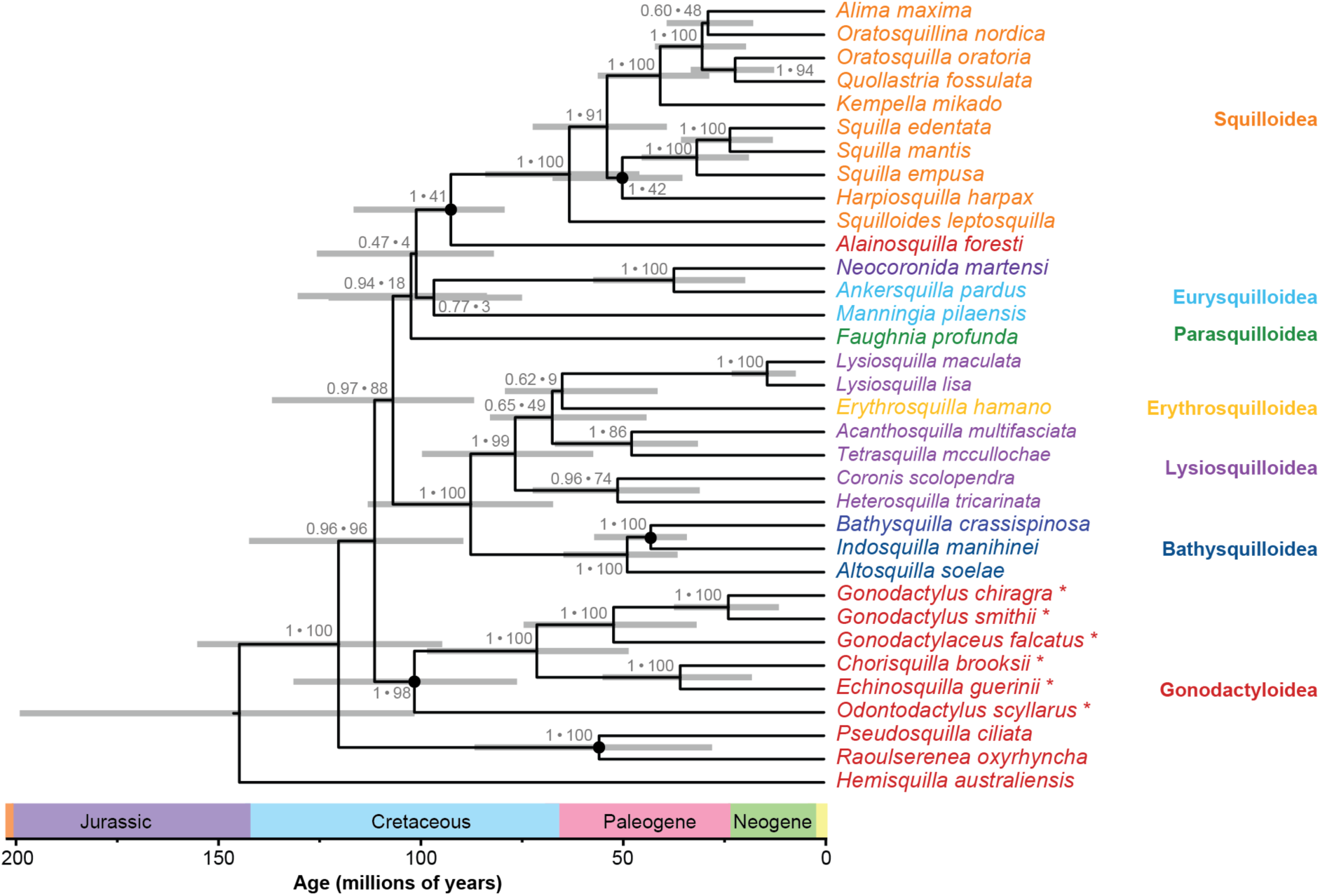
Total-evidence Bayesian phylogenetic estimate of Stomatopoda, inferred from 77 morphological characters and 14,598 nucleotide sites. Branch lengths are proportional to time. Taxon colors correspond to their superfamily classification. Support values at nodes correspond to posterior probability (>0.95) and likelihood bootstrap support (>75%). Calibrated nodes are represented by black circles. Horizontal grey bars denote 95% credibility intervals of node-age estimates. Asterisks indicate species with specialized ‘smashing’ appendages.

The Bayesian estimates of divergence times were robust to the choice of tree prior, while comparison of marginal likelihoods showed very strong support for a Yule model over a birth-death model (2 log Bayes factor > 10). Our molecular dating analyses provided an posterior median estimate of 140 Ma (95% credible interval 201–102 Ma) for the age of crown-group unipeltatan stomatopods. The date estimates suggest that all of the stomatopod superfamilies had diverged from each other by the beginning of the Paleogene (66 Ma). The divergence between smashing and spearing stomatopods occurred 98 Ma (95% CI 131–76 Ma).

We found no evidence of correlation between rates of morphological and molecular evolution across the branches of the phylogeny. This result was obtained for the combined molecular clock (*p* = 0.184; Fig. 4A) and when the mitochondrial and nuclear genes were assigned separate clock models (Fig. 4B). We did not detect a correlation between morphological and molecular rates after excluding terminal branches, on which morphological rates might be underestimated because of undersampling of autapomorphies (Seligmann, 2010; Lee et al., 2013). The degree of among-lineage rate heterogeneity, as measured by the coefficient of variation of rates (Drummond et al., 2006), was much higher for the morphological data (3.27, 95% CI 2.12–4.66) than for the molecular data (0.39, 95% CI 0.30–0.48) (Fig. 5).

**Fig. 4.**
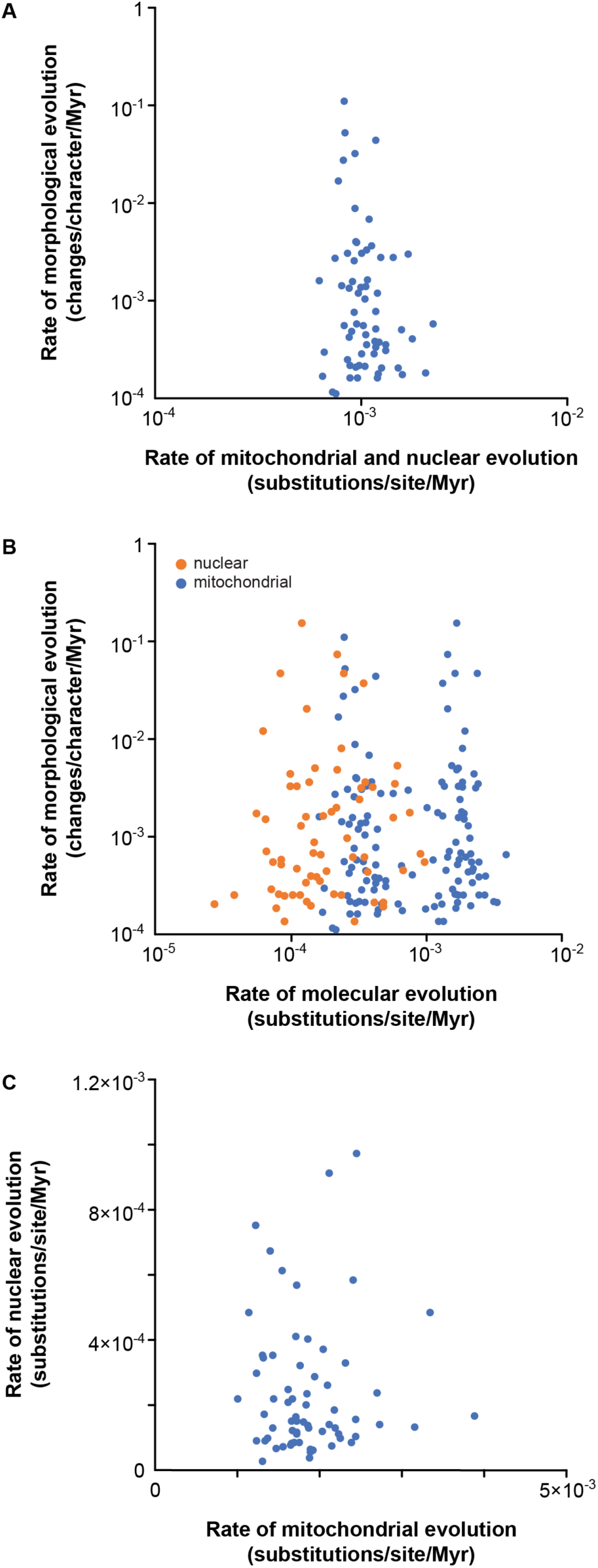
Rates of morphological and molecular evolution for each branch in the inferred stomatopod phylogeny, with (A) mitochondrial and nuclear data combined or (B) mitochondrial and nuclear evolutionary rates estimated separately. (C) Comparison of rates of nuclear and mitochondrial evolution for each branch.

**Fig. 5.**
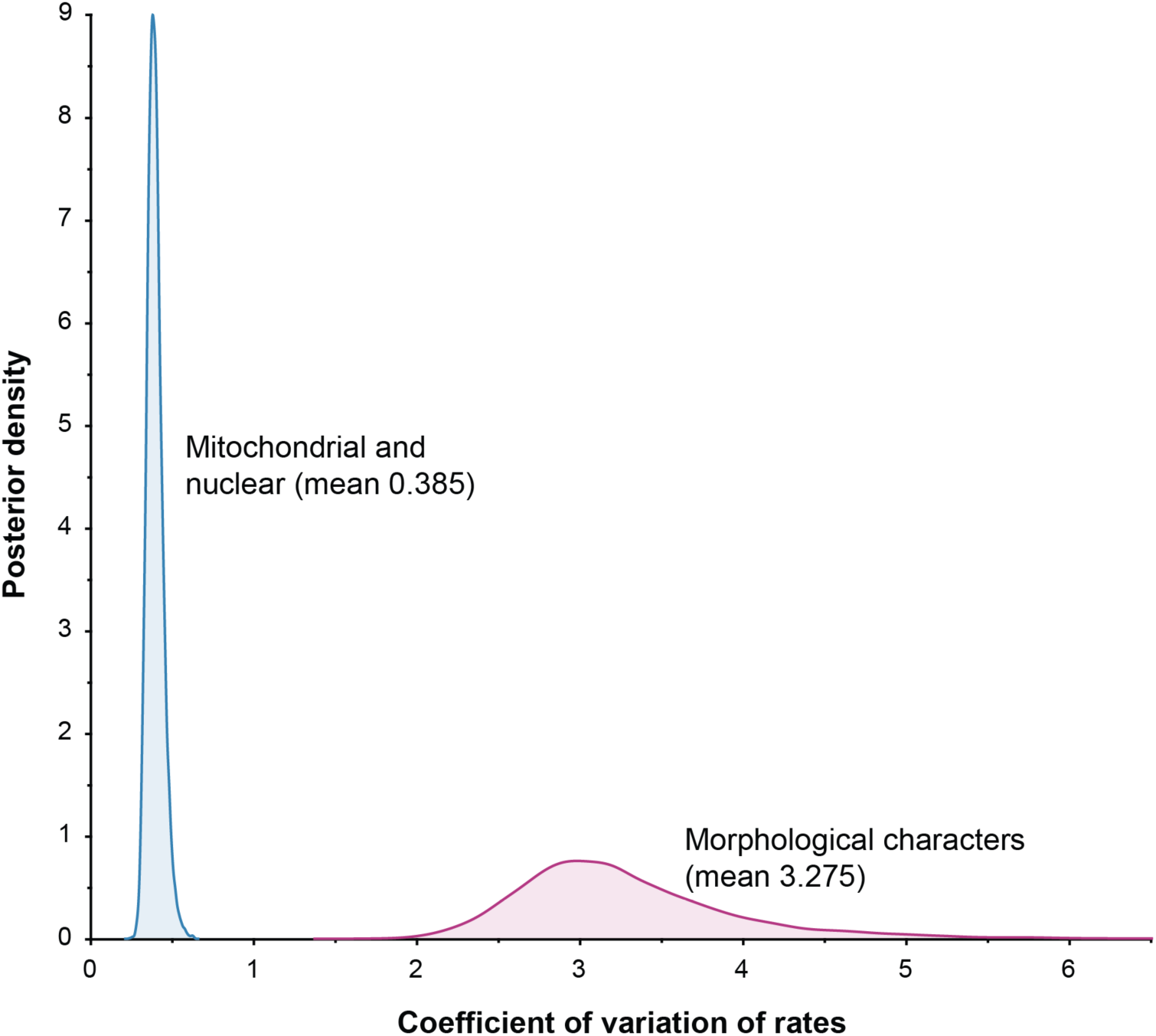
Coefficient of variation of stomatopod evolutionary rates in molecular and morphological characters, estimated in a Bayesian relaxed-clock analysis of a combined molecular and morphological data set.

Our Bayesian phylogenetic analyses yielded no evidence of correlation between rates of mitochondrial and nuclear evolution across the branches of the phylogeny (*p* = 0.929; Fig. 4C). As expected, substitution rates were lower in nuclear genes (mean rate 2.25×10^-4^, 95% CI 1.58×10^-4^–2.85×10^-4^ substitutions per site per Myr) than in mitochondrial genes (mean rate 1.80×10^-3^, 95% CI 1.29×10^-3^–2.26×10^-3^ substitutions per site per Myr). The coefficient of variation of rates was higher for the nuclear markers (1.03, 95% CI 0.80–1.27) than for the mitochondrial genomes (0.38, 95% CI 0.29–0.47).

## 4. Discussion

### 4.1. Phylogenetic relationships

This study has provided a total-evidence phylogenetic estimate for the seven recognized superfamilies of Stomatopoda. Our estimate of the stomatopod phylogeny based on combined morphological and molecular data is better supported than the trees inferred using the two data types separately. The results of our analyses underscore previous conclusions about the limitations of mitochondrial genomes in resolving deep stomatopod relationships (Koga and Rouse, 2021). Here we have recovered evolutionary signals from both data sets, revealing relationships that were present in the separate morphological and molecular trees. A similar improvement in nodal support has also been seen in other studies of combined morphological and molecular data, particularly in cases where molecular sequence data were incomplete for some taxa (Hillis, 1987; Lee et al., 2007; Bracken-Grissom et al., 2013; Ruhfel et al., 2013; Bracken-Grissom et al., 2014).

Our total-evidence phylogeny of Stomatopoda is congruent with previous evidence of a sister relationship between major clades containing Lysiosquilloidea and Squilloidea (Ahyong and Harling, 2000; Ahyong and Jarman, 2009). This result is in contrast with some previous findings of a sister relationship between Gonodactyloidea and Squilloidea (Porter et al., 2010; Van Der Wal et al., 2017; Koga and Rouse, 2021). The placement of Bathysquilloidea in our phylogenetic estimate also contrasts with that inferred by Van Der Wal et al. (2017). Here we find strong support for Bathysquilloidea being closely related to Lysiosquilloidea, rather than to Gonodactyloidea (Van Der Wal et al., 2017) or to a clade comprising Squilloidea, Parasquilloidea, and Eurysquilloidea (Ahyong and Harling, 2000). The close relationship between Eurysquilloidea, Parasquilloidea, and Squilloidea identified by Van Der Wal et al. (2017) is also recovered here, and is consistent with previous morphological estimates (Ahyong and Harling, 2000; Ahyong, 2001, 2005).

The placement of the coronidid lysiosquilloid, *Neocoronida martensi* Manning, 1978, within Eurysquilloidea is noteworthy. The elaborate dorsal telson ornamentation and aspects of the raptorial claw of *Neocoronida* led to its original placement in the lysiosquilloids, but other features of telson denticulation and uropod and maxillipedal structure align with those of the eurysquilloids, especially *Ankersquilla* and *Liusquilla* (Ahyong and Lin, 2020; Ahyong et al., 2020). Morphological re-evaluations currently underway, together with our phylogenetic results, indicate that *Neocoronida* should be transferred from Lysiosquilloidea to Eurysquilloidea.

The lack of support for a monophyletic Gonodactyloidea indicates that revisions within this superfamily are required (Koga and Rouse, 2021). Specifically, our estimated placement of Pseudosquillidae corroborates the findings of previous molecular studies, suggesting that pseudosquillids are not true gonodactyloids and should be moved to a new superfamily (Ahyong and Jarman, 2009; Porter et al., 2010; Koga and Rouse, 2021). Similarly, the inferred position of the morphologically intermediate Alainosquillidae differs from morphological estimates (Ahyong and Harling, 2000), indicating that this family might not be a true gonodactyloid; it may require reclassification and is currently the subject of further study. The position of Odontodactylidae as the sister group to all other specialized smashing stomatopods is consistent with previous morphological and molecular analyses of Gonodactyloidea (Ahyong and Harling, 2000; Barber and Erdmann, 2000; Porter et al., 2010), which also inferred a monophyletic group of smashing stomatopods.

Our study presents the first sequence data from any member of the superfamily Erythrosquilloidea, represented by *Erythrosquilla hamano* Ahyong, 2001. *Erythrosquilla* and the lysiosquilloids form a strongly supported clade that corroborates the morphological synapomorphies uniting the two groups, particularly the ventrally ribbed maxilliped 3–4 propodi (Ahyong and Harling, 2000). In our inferred phylogenies, however, *Erythrosquilla hamano* was consistently nested within, rather than placed as a sister lineage to, Lysiosquilloidea, albeit with only moderate support. This result does not support the separate superfamily status of Erythrosquilloidea, but is nevertheless consistent with morphological evidence of a close relationship between Erythrosquilloidea and Lysiosquilloidea (Ahyong, 1997; Ahyong and Harling, 2000). Ahyong (2001) previously noted that *Erythrosquilla* resembles a number of lysiosquilloid genera in telson and uropod morphology. Further phylogenetic study is required to corroborate the present results and to determine whether Erythrosquilloidea should continue to be recognized as a separate superfamily or subsumed within Lysiosquilloidea.

### 4.2. Evolutionary timescale of Stomatopoda

Our estimate of the stomatopod evolutionary timescale indicates that diversification events within the group occurred more recently than previously inferred. We estimate that crown-group unipeltatan stomatopods appeared ∼140 Ma in the late Mesozoic. The posterior median estimate is approximately 50 Myr younger than in previous analyses, but lies within the reported 95% CI of the estimate by Van Der Wal et al. (2017). The divergences among superfamilies appear to have taken place well before the separation of the Atlantic and Pacific Oceans ∼35 Ma in the Eocene, which is consistent with the worldwide distributions of the major stomatopod lineages (Ekman, 1953; Reaka et al., 2008).

Our results agree with those of Van Der Wal et al. (2017) in supporting the hypothesis that the formation of new coastal habitat during the breakup of Pangaea and Gondwana allowed stomatopods to diversify rapidly and to expand their range. These biogeographic events shaped the distribution patterns of many marine groups, including anomuran and brachyuran crabs (Bracken-Grissom et al., 2013; Tsang et al., 2014), underpinning the Tethyan patterns of distribution that are seen today (George, 2006).

The divergence between smashing and spearing stomatopods in the Mesozoic supports the hypothesis that specialized smashing and spearing appendages appeared early in the evolutionary history of Unipeltata (Ahyong and Harling, 2000; Ahyong and Jarman, 2009). This contrasts with the hypothesis that smashing stomatopods evolved after a long history of spearing (Caldwell, 1991; Ahyong, 1997; Koga and Rouse, 2021). Our results indicate that the two monophyletic groups most likely diverged from an unspecialized or intermediate stomatopod ancestor, perhaps similar to present-day *Hemisquilla* (Ahyong and Harling, 2000; Ahyong and Jarman, 2009; Braig et al., 2023).

### 4.3. Rates of morphological and molecular evolution

We did not find evidence of a correlation between rates of morphological and molecular evolution across branches of the stomatopod tree. These results support the findings of previous studies of morphological and molecular evolutionary rates in hermit, king, and horseshoe crabs (Cunningham et al., 1992; Avise et al., 1994), as well as broader studies of evolutionary rates in animals (Davies and Savolainen, 2006; Halliday et al., 2019) and flowering plants (Asar et al., 2023). The decoupling of morphological and molecular rates might be attributed to different adaptive constraints acting on these data sets (Lee and Palci, 2015). Alternatively, we might not be able to detect a correlation in these rates if very little of the genome is linked directly to phenotypic and adaptive changes (Bromham et al., 2002), or if some morphological variations are non-heritable (Hillis, 1987).

The substantial differences in evolutionary rate variation among lineages can partly explain the lack of a correlation between morphological and molecular rates. In agreement with previous studies, we found much greater among-lineage rate variation in morphological characters than in the molecular data (Beck and Lee, 2014; Harrison and Larsson, 2014). Further insights into the causes of such rate variation in morphological data sets might come from comparing different partitioning schemes, such as by anatomical area, position on the body, or type of character (Lee, 2016; Simões and Pierce, 2021). This would help to elucidate whether the evolution of these characters is subject to a ‘common mechanism’ (Goloboff et al., 2019). In this context, the tools designed for partitioning molecular clocks might be useful for comparing different clock-partitioning schemes for morphological characters (Duchêne et al., 2014; Lee, 2016; Duchêne et al., 2024).

The choice of data sets in our analysis might also have some bearing on the lack of a correlation between molecular and morphological evolutionary rates. Analysing larger numbers of morphological characters will increase the power to detect a correlation with rates of molecular evolution (Omland, 1997; Asar et al., 2023). For example, Lee et al. (2013) found evidence of a correlation between morphological and molecular rates in arthropods during the Cambrian explosion, using a large data set that comprised 395 morphological characters and 62 protein-coding genes. Our morphological characters might not be sufficiently granular to capture the overall rate of morphological evolution in stomatopod lineages, because many were scored for their effectiveness across relatively high taxonomic levels. Alternatively, our molecular data set might not include the small proportion of genes in the genome that are expected to be associated with measurable phenotypic changes. This possibility is reflected in the lack of a correlation between the rates in mitochondrial genomes and the nuclear genes in our data set, a finding that is consistent with previous evidence from metazoans (Lynch et al., 2006; Allio et al., 2017). A data set comprising a large number of nuclear genes with denser taxon sampling, along with an expanded set of morphological characters, is likely to improve our power to detect any rate correlations.

## 5. Conclusions

We have used a total-evidence approach to obtain a robust estimate of the stomatopod phylogeny, including the relationships among all seven extant stomatopod superfamilies. Our analysis of morphological and molecular rates did not yield evidence of a correlation, but did indicate very high levels of among-lineage rate variation in the morphological data set. The decoupling of morphological and molecular rates within Stomatopoda is consistent with the results of previous crustacean studies. Our study has highlighted the merit of combining morphological and molecular data in evolutionary analyses. Future studies using genome-wide data and increased taxon sampling will improve resolution of the phylogenetic relationships and evolutionary rates of stomatopods.

## Acknowledgements

We acknowledge funding from a Graduate Student Research Award from the Society of Systematic Biologists and a Joyce W. Vickery Scientific Research Fund grant from the Linnean Society to CVDW. We thank Michael Lee from the South Australian Museum for advice on data analyses. This research was partly supported by Discovery Project DP220103265 from the Australian Research Council to NL and SYWH. STA gratefully acknowledges partial support from the Singapore Ministry of Defence and Lee Kong Chian Natural History Museum (National University of Singapore). The authors acknowledge the Sydney Informatics Hub and the University of Sydney’s high-performance computing cluster Artemis for providing the computing resources that have contributed to the research results reported within this paper.

## References

Ahyong, S.T., 1997. Phylogenetic analysis of the Stomatopoda (Malacostraca). J. Crustacean Biol. 17, 695–715.

Ahyong, S.T., 2001. Revision of the Australian Stomatopod Crustacea. Rec. Aust. Mus. Suppl. 26, 1–326.

Ahyong, S.T., 2005. Phylogenetic analysis of the Squilloidea (Crustacea : Stomatopoda). Invertebr. Syst. 19, 189–208.

Ahyong, S.T., Harling, C., 2000. The phylogeny of the stomatopod Crustacea. Aust. J. Zool. 48, 607–642.

Ahyong, S.T., Jarman, S.N., 2009. Stomatopod interrelationships: Preliminary results based on analysis of three molecular loci. Arthropod Syst. Phylo. 67, 91–98.

Ahyong, S.T., Lin, C.-W., 2020. The mantis shrimp superfamily Eurysquilloidea confirmed from Taiwan: *Liusquilla formosa* gen. et sp. nov. Crustaceana 93, 1473–1482.

Ahyong, S.T., Porter, M.L., Caldwell, R.L., 2020. The Leopard Mantis Shrimp, *Ankersquilla pardus*, a new genus and species of eurysquillid from Indo-West Pacific coral reefs. Rec. Aust. Mus. 72, 1–8.

Allio, R., Donega, S., Galtier, N., Nabholz, B., 2017. Large variation in the ratio of mitochondrial to nuclear mutation rate across animals: Implications for genetic diversity and the use of mitochondrial DNA as a molecular marker. Mol. Biol. Evol. 34, 2762–2772.

Asar, Y., Sauquet, H., Ho, S.Y.W., 2023. Evaluating the accuracy of methods for detecting correlated rates of molecular and morphological evolution. Syst. Biol., 72, 1337– 1356.

Avise, J.C., Nelson, W.S., Sugita, H., 1994. A speciational history of “living fossils”: Molecular evolutionary patterns in horseshoe crabs. Evolution 48, 1986–2001.

Barber, P.H., Erdmann, M.V., 2000. Molecular systematics of the Gonodactylidae (Stomatopoda) using mitochondrial cytochrome oxidase c (subunit 1) DNA sequence data. J. Crustacean Biol. 20, 20–36.

Beck, R.M., Lee, M.S.Y., 2014. Ancient dates or accelerated rates? Morphological clocks and the antiquity of placental mammals. Proc. Biol. Sci. 281, e20141278.

Bernt, M., Donath, A., Juhling, F., Externbrink, F., Florentz, C., Fritzsch, G., Putz, J., Middendorf, M., Stadler, P.F., 2013. MITOS: Improved *de novo* metazoan mitochondrial genome annotation. Mol. Phylogenet. Evol. 69, 313–319.

Bouckaert, R., Vaughan, T.G., Barido-Sottani, J., Duchêne, S., Fourment, M., Gavryushkina, A., Heled, J., Jones, G., Kühnert, D., De Maio, N., Matschiner, M., Mendes, F.K., Müller, N.F., Ogilvie, H.A., du Plessis, L., Popinga, A., Rambaut, A., Rasmussen, D., Siveroni, I., Suchard, M.A., Wu, C.-H., Xie, D., Zhang, C., Stadler, T., Drummond, A.J., 2019. BEAST 2.5: An advanced software platform for Bayesian evolutionary analysis. PLOS Comput. Biol. 15, e1006650.

Bouckaert, R.R., Drummond, A.J., 2017. bModelTest: Bayesian phylogenetic site model averaging and model comparison. BMC Evol. Biol. 17, 42.

Bracken-Grissom, H.D., Ahyong, S.T., Wilkinson, R.D., Feldmann, R.M., Schweitzer, C.E., Breinholt, J.W., Bendall, M., Palero, F., Chan, T.-Y., Felder, D.L., Robles, R., Chu, K.-H., Tsang, L.-M., Kim, D., Martin, J.W., Crandall, K.A., 2014. The emergence of lobsters: Phylogenetic relationships, morphological evolution and divergence time comparisons of an ancient group (Decapoda: Achelata, Astacidea, Glypheidea, Polychelida). Syst. Biol. 63, 457–479.

Bracken-Grissom, H.D., Cannon, M.E., Cabezas, P., Feldmann, R.M., Schweitzer, C.E., Ahyong, S.T., Felder, D.L., Lemaitre, R., Crandall, K.A., 2013. A comprehensive and integrative reconstruction of evolutionary history for Anomura (Crustacea: Decapoda). BMC Evol. Biol. 13, 128.

Braig, F., Haug, J. T., Ahyong, S. T., Garassino, A., Schädel, M., Haug, C., 2023. Another piece in the puzzle of mantis shrimp evolution – fossils from the Early Jurassic Osteno Lagerstätte of Northern Italy. Compt. Rend. Palevol 22, 17–31.

Bromham, L., Woolfit, M., Lee, M.S.Y., Rambaut, A., 2002. Testing the relationship between morphological and molecular rates of change along phylogenies. Evolution 56, 1921– 1930.

Caldwell, R.L., 1991. Stomatopods: the better to see you with my dear. Aust. Nat. Hist. 23, 696–705.

Cunningham, C.W., Blackstone, N.W., Buss, L.W., 1992. Evolution of king crabs from hermit crab ancestors. Nature 355, 539–542.

Davies, T.J., Savolainen, V., 2006. Neutral theory, phylogenies, and the relationship between phenotypic change and evolutionary rates. Evolution 60, 476–483.

Drummond, A.J., Ho, S.Y.W., Phillips, M.J., Rambaut, A., 2006. Relaxed phylogenetics and dating with confidence. PLOS Biol. 4, e88.

Duchêne, D.A., Duchêne, S., Ho, S.Y.W., 2017. New statistical criteria detect phylogenetic bias caused by compositional heterogeneity. Mol. Biol. Evol. 34, 1529–1534.

Duchêne, D.A., Duchêne, S., Ho, S.Y.W., 2018. PhyloMAd: efficient assessment of phylogenomic model adequacy. Bioinformatics 34, 2300–2301.

Duchêne, D.A., Duchêne, S., Stiller, J., Heller, R., Ho, S.Y.W., 2024. ClockstaRX: Testing molecular clock hypotheses with genomic data. Genome Biol. Evol. 16, evae064.

Duchêne, S., Molak, M., Ho, S.Y.W., 2014. ClockstaR: choosing the number of relaxed-clock models in molecular phylogenetic analysis. Bioinformatics 30, 1017–1019.

Duchêne, S., Tong, K.J., Foster, C.S.P., Duchêne, S., Lanfear, R., Ho, S.Y.W., 2020. Linking branch lengths across sets of loci provides the highest statistical support for phylogentic inference. Mol Biol Evol 37, 1202–1210.

Edgar, R.C., 2004. MUSCLE: multiple sequence alignment with high accuracy and high throughput. Nucleic Acids Res. 32, 1792–1797.

Ekman, S., 1953. Zoogeography of the sea. Sidgwick & Jackson, London. 417 pp.

Franklin, A.M., Donatelli, C.M., Culligan, C.R., Tytell, E.D., 2019. Meral-spot reflectance signals weapon performance in the mantis shrimp *Neogonodactylus oerstedii* (Stomatopoda). Biol. Bull. 236, 43–54.

Gagnon, Y.L., Templin, R.M., How, M.J., Marshall, N.J., 2015. Circularly polarized light as a communication signal in mantis shrimps. Curr. Biol. 25, 3074–3078.

George, R.W., 2006. Tethys origin and subsequent radiation of the spiny lobsters (Palinuridae). Crustaceana 79, 397–422.

Goloboff, P.A., Pittman, M., Pol, D., Xu, X., 2019. Morphological data sets fit a common mechanism much more poorly than DNA sequences and call into question the Mkv model. Syst. Biol. 68, 494–504.

Green, P.A., Patek, S.N., 2018. Mutual assessment during ritualized fighting in mantis shrimp (Stomatopoda). Proc. R. Soc. B 285, 20172542.

Halliday, T.J.D., dos Reis, M., Tamuri, A.U., Ferguson-Gow, H., Yang, Z., Goswami, A., 2019. Rapid morphological evolution in placental mammals post-dates the origin of the crown group. Proc. R. Soc. B 286, 20182418.

Harrison, L.B., Larsson, H.C.E., 2014. Among-character rate variation distributions in phylogenetic analysis of discrete morphological characters. Syst. Biol. 64, 307–324.

Haug, C., Kutschera, V., Ahyong, S. T., Vega, F. J., Maas, A., Waloszek, D., Haug, J. T., 2013. Re-evaluation of the Mesozoic mantis shrimp *Ursquilla yehoachi* based on new material and the virtual peel technique. Palaeontol. Electron., 16, 5T1–14.

Hillis, D.M., 1987. Molecular versus morphological approaches to systematics. Annu. Rev. Ecol. Syst. 18, 23–42.

Hwang, H.-S., Jung, J., 2021. First record of the complete mitochondrial genome of the mantis shrmp, *Gonodactylaceus randalli* (Manning, 1978) (Stomatopoda: Gonodactylidae). Mitochondrial DNA B 6, 510–511.

Hwang, H.-S., Jung, J., Baeza, J.A., 2021. The mitochondrial genome of *Faughnia haani* (Stomatopoda): novel organization of the control region and phylogenetic position of the superfamily Parasquilloidea. BMC Genomics 22, 716.

Kass, R.E., Raftery, A.E., 1995. Bayes factors. J. Am. Stat. Assoc. 90, 773–795.

Koga, C., Rouse, G.W., 2021. Mitogenomics and the phylogeny of mantis shrimps (Crustacea: Stomatopoda). Diversity 13, 647.

Lanfear, R., Calcott, B., Ho, S.Y.W., Guindon, S., 2012. PartitionFinder: Combined selection of partitioning schemes and substitution models for phylogenetic analyses. Mol. Biol. Evol. 29, 1695–1701.

Lanfear, R., Frandsen, P.B., Wright, A.M., Senfeld, T., Calcott, B., 2017. PartitionFinder 2: New methods for selecting partitioned models of evolution for molecular and morphological phylogenetic analyses. Mol. Biol. Evol. 34, 772–773.

Lee, M.S.Y., 2016. Multiple morphological clocks and total-evidence tip-dating in mammals. Biol. Lett. 12, 20160033.

Lee, M.S.Y., Hugall, A.F., Lawson, R., Scanlon, J.D., 2007. Phylogeny of snakes (Serpentes): combining morphological and molecular data in likelihood, Bayesian and parsimony analyses. Syst. Biodivers. 5, 371–389.

Lee, M.S.Y., Palci, A., 2015. Morphological phylogenetics in the genomic age. Curr. Biol. 25, R922–R929.

Lee, M.S.Y., Soubrier, J., Edgecombe, G.D., 2013. Rates of phenotypic and genomic evolution during the Cambrian explosion. Curr. Biol. 23, 1889–1895.

Lewis, P.O., 2001. A likelihood approach to estimating phylogeny from discrete morphological character data. Syst. Biol. 50, 913–925.

Liu, Y.A., Cui, Z.X., 2010. The complete mitochondrial genome of the mantid shrimp *Oratosquilla oratoria* (Crustacea: Malacostraca: Stomatopoda): Novel non-coding regions features and phylogenetic implications of the Stomatopoda. Comp. Biochem. Phys. D 5, 190–198.

Lynch, M., Koskella, B., Schaack, S., 2006. Mutation pressure and the evolution of organelle genomic architecture. Science 311, 1727–1730.

Manning, R.B., 1968. A revision of the family Squillidae (Crustacea, Stomatopoda), with the description of eight new genera. Bull. Mar. Sci. 18, 105–142.

Manning, R. B., 1978. New and rare Stomatopod Crustacea from the Indo-West Pacific region. Smithson. Contrib. Zool. 264, 1–36.

Manning, R.B., 1980. The superfamilies, families, and genera of Recent Stomatopod Crustacea, with diagnoses of six new families. Proc. Biol. Soc. Wash. 93, 362–372.

Marshall, J., Cronin, T.W., Kleinlogel, S., 2007. Stomatopod eye structure and function: A review. Arthropod Struct. Dev. 36, 420–448.

Nguyen, J.M.T., Ho, S.Y.W., 2020. Calibrations from the fossil record. In: Ho, S.Y.W. (ed) The Molecular Evolutionary Clock: Theory and Practice. Springer, Cham, Switzerland. pp. 117–133.

Omland, K.E., 1997. Correlated rates of molecular and morphological evolution. Evolution 51, 1381–1393.

Patek, S.N., 2019. The power of extreme movement: evolution, behavior, and biomechanics of mantis shrimp strikes. Integr. Comp. Biol. 59, e180.

Porter, M.L., Zhang, Y.F., Desai, S., Caldwell, R.L., Cronin, T.W., 2010. Evolution of anatomical and physiological specialization in the compound eyes of stomatopod crustaceans. J. Exp. Biol. 213, 3473–3486.

Rambaut, A., Drummond, A.J., Xie, D., Baele, G., Suchard, M.A., 2018. Posterior summarization in Bayesian phylogenetics using Tracer 1.7. Syst. Biol. 67, 901–904.

Reaka, M.L., Manning, R.B., 1981. The behavior of stomatopod Crustacea, and its relationship to rates of evolution. J. Crustacean Biol. 1, 309–327.

Reaka, M.L., Manning, R.B., 1987. The significance of body size, dispersal potential, and habitat for rates of morphological evolution in stomatopod Crustacea. Smithsonian Institution Press, Washington, DC, USA. 46 pp.

Reaka, M.L., Rodgers, P.J., Kudla, A.U., 2008. Patterns of biodiversity and endemism on Indo-West Pacific coral reefs. Proc. Natl Acad. Sci. USA 105, 11474–11481.

Ronquist, F., Teslenko, M., van der Mark, P., Ayres, D.L., Darling, A., Höhna, S., Larget, B., Liu, L., Suchard, M.A., Huelsenbeck, J.P., 2012. MrBayes 3.2: Efficient Bayesian phylogenetic inference and model choice across a large model space. Syst. Biol. 61, 539–542.

Ruhfel, B.R., Stevens, P.F., Davis, C.C., 2013. Combined morphological and molecular phylogeny of the clusioid clade (Malpighiales) and the placement of the ancient rosid macrofossil *Paleoclusia*. Int. J. Plant Sci. 174, 910–936.

Schram, F.R., 2007. Paleozoic proto-mantis shrimp revisited. J. Paleontol. 81, 895–916.

Schram, F.R., Klein, J.C.v.V., Charmantier-Daures, M., 2013. Treatise on Zoology - Anatomy, Taxonomy, Biology. The Crustacea, Volume 4 Part A. Brill, Leiden, The Netherlands. 124 pp.

Seligmann, H., 2010. Positive correlations between molecular and morphological rates of evolution. J. Theor. Biol. 264, 799–807.

Simões, T.R., Pierce, S.E., 2021. Sustained high rates of morphological evolution during the rise of tetrapods. Nat. Ecol. Evol. 5, 1403–1414.

Stadler, T., 2009. On incomplete sampling under birth-death models and connections to the sampling-based coalescent. J. Theor. Biol. 261, 58–66.

Stamatakis, A., 2014. RAxML version 8: a tool for phylogenetic analysis and post-analysis of large phylogenies. Bioinformatics 30, 1312–1313.

Thomas, J.A., Welch, J.J., Woolfit, M., Bromham, L., 2006. There is no universal molecular clock for invertebrates, but rate variation does not scale with body size. Proc. Natl Acad. Sci. USA 103, 7366–7371.

Tsang, L.M., Schubart, C.D., Ahyong, S.T., Lai, J.C., Au, E.Y., Chan, T.Y., Ng, P.K., Chu, K.H., 2014. Evolutionary history of true crabs (Crustacea: Decapoda: Brachyura) and the origin of freshwater crabs. Mol. Biol. Evol. 31, 1173–1187.

Van Der Wal, C., Ahyong, S.T., Ho, S.Y.W., Lo, N., 2017. The evolutionary history of Stomatopoda (Crustacea: Malacostraca) inferred from molecular data. PeerJ 5, e3844.

Wolfe, J.M., Daley, A.C., Legg, D.A., Edgecombe, G.D., 2016. Fossil calibrations for the arthropod Tree of Life. Earth-Sci. Rev. 160, 43–110.

Xia, X., 2017. DAMBE6: New tools for microbial genomics, phylogenetics, and molecular evolution. J. Hered. 108, 431–437.

Xie, W.G., Lewis, P.O., Fan, Y., Kuo, L., Chen, M.H., 2011. Improving marginal likelihood estimation for Bayesian phylogenetic model selection. Syst. Biol. 60, 150–160.

Yang, M., Liu, H., Wang, R., Tan, W., 2021. The complete mitochondrial genome of Purple Spot Mantis Shrimp *Gonodactylus smithii* (Pocock, 1893). Mitochondrial DNA B 6, 2028–2030.

