## Supplementary Material for "Total-evidence phylogenetic analysis resolves the evolutionary timescale of mantis shrimps (Stomatopoda) and provides insights into their molecular and morphological evolutionary rates"

**Supplementary Table S1.** List of stomatopod species analysed in this study, with corresponding GenBank accession numbers. **BOLD** accessions denote new sequences generated in this study, dashes (-) indicate missing sequences, asterisks (*) indicate incomplete mitogenomes for which gene-specific accession numbers are provided in Supplementary Table S2.

| **Taxon** | **Mitochondrial genome** | ***18S*** | ***28S*** | ***H3*** |
|---|---|---|---|---|
| **Bathysquilloidea** | | | | |
| *Altosquilla soelae* (Bruce, 1985) | * | **MN306278** | **MN338474** | **MN329775** |
| *Bathysquilla crassispinosa* (Fukuda, 1909) | **PP761313** | **MN306284** | **MN338473** | **MN329774** |
| *Indosquilla manihinei* (Ingle & Merrett, 1971) | * | **MN306280** | **MN338475** | **MN329773** |
| **Erythrosquilloidea** | | | | |
| *Erythrosquilla hamano* (Ahyong, 2001) | * | – | **MN338476** | – |
| **Eurysquilloidea** | | | | |
| *Ankersquilla pardus* (Ahyong, Porter & Caldwell, 2020) | **PP761316** | **OR775461** | **OR781484, OR781485** | **OR782937** |
| *Manningia pilaensis* (de Man, 1888) | * | **MN306274** | **MN338461** | **MN329770** |
| **Gonodactyloidea** | | | | |
| *Alainosquilla foresti* (Moosa, 1991) | * | **MN306279** | **MN338467** | **MN329771** |
| *Chorisquilla brooksii* (de Man, 1888) | **PP761320** | **MN306264** | **MN338456** | **MN329764** |
| *Echinosquilla guerinii* (White, 1861) | * | **MN306265** | **MN338455** | **MN329763** |
| *Gonodactylaceus falcatus* (Forskål, 1775) | **PP740381** | HM138871 | HM180015 | – |
| *Gonodactylus chiragra* (Fabricius, 1781) | DQ191682 | HM138870 | HM180014 | – |
| *Gonodactylus smithii* (Pocock, 1893) | * | **MN306273** | **MN338454** | **MN329755** |
| *Hemisquilla australiensis* (Stephenson, 1967) | **PP761312** | **MN306283** | **MN338464** | **MN329753** |
| *Odontodactylus scyllarus* (Linnaeus, 1758) | **PP761321** | **MN306271** | **MN338466** | **MN329769** |
| *Pseudosquilla ciliata* (Fabricius, 1787) | AY947836 | HM138888 | HM180032 | – |

| | | | | |
|---|---|---|---|---|
| *Raoulserenea oxyrhyncha* (Borradaile, 1898) | **PP761322** | **MN306275** | **MN338463** | **MN329754** |
| **Lysiosquilloidea** | | | | |
| *Acanthosquilla multifasciata* (Wood-Mason, 1895) | * | **MN306262** | **MN338458** | **MN329765** |
| *Coronis scolopendra* (Latreille, 1828) | **PP761317** | HM138863 | HM180007 | – |
| *Heterosquilla tricarinata* (Claus, 1871) | * | **MN306266** | **MN338471** | **MN329772** |
| *Lysiosquilla lisa* (Ahyong & Randall, 2001) | * | **MN306277** | **MN338462** | **MN329762** |
| *Lysiosquilla maculata* (Fabricius, 1793) | DQ191683 | HM138878 | EU920998 | EU921076 |
| *Neocoronida martensi* (Manning, 1978) | **PP761315** | **MN306272** | **MN338459** | **MN329767** |
| *Tetrasquilla mccullochae* (Schmitt, 1940) | * | **MN306276** | **MN338472** | **MN329766** |
| **Parasquilloidea** | | | | |
| *Faughnia profunda* (Manning & Makarov, 1978) | PP761314 | **MN306282** | **MN338457** | **MN329768** |
| **Squilloidea** | | | | |
| *Alima maxima* (Ahyong, 2002) | * | **MN306267** | **MN338468** | **MN329757** |
| *Harpiosquilla harpax* (de Haan, 1844) | AY699271 | – | FJ871143 | – |
| *Kempella mikado* (Kemp & Chopra, 1921) | **PP761318** | **MN306268** | **MN338465** | **MN329758** |
| *Oratosquilla oratoria* (de Haan, 1844) | GQ292769 | | LC167473 | **MN329761** |
| *Oratosquillina nordica* (Ahyong & Chan, 2008) | * | **MN306263** | **MN338470** | **MN329760** |
| *Quollastria fossulata* (Moosa, 1986) | **PP761319** | **MN306270** | **MN338469** | **MN329756** |
| *Squilla edentata* (Lunz, 1937) | * | **MN306269** | **MN338460** | **MN329759** |
| *Squilla empusa* (Say, 1818) | DQ191684 | L81946 | AY210842 | JN800718 |
| *Squilla mantis* (Linnaeus, 1758) | AY639936 | GQ328958 | – | – |
| *Squilloides leptosquilla* (Brooks, 1886) | KR095170 | – | MH168185 | – |

**Supplementary table S2.** List of stomatopod species for which incomplete mitochondrial genomes were generated in this study, with corresponding gene-specific GenBank accession numbers. Dashes (-) indicate missing sequences.

| Taxon | *ATP6* | *ATP8* | *COB* | *COX1* | *COX2* | *COX3* | *NAD1* | *NAD2* | *NAD3* | *NAD4* | *NAD4L* | *NAD5* | *NAD6* | *12S* | *16S* |
|---|---|---|---|---|---|---|---|---|---|---|---|---|---|---|---|
| **Bathysquilloidea** | | | | | | | | | | | | | | | |
| *Altosquilla soelae* | PP785047 | PP785057 | PP768095 | PP796532 | PP768008 | PP768018 | PP768029 | PP785066 | PP785080 | PP768041 | - | - | PP785100 | PP768071 | PP768083 |
| *Indosquilla manihinei* | - | - | PP795577 | - | - | PP795579 | - | - | - | - | - | - | - | - | - |
| **Erythrosquilloidea** | | | | | | | | | | | | | | | |
| *Erythrosquilla hamano* | - | - | - | - | - | PP795578 | - | - | PP795576 | - | - | - | - | - | - |
| **Eurysquilloidea** | | | | | | | | | | | | | | | |
| *Manningia pilaensis* | PP785040 | PP785052 | PP768089 | PP796528 | PP768001 | PP768013 | - | PP785062 | PP785073 | PP768035 | PP785085 | PP768047 | PP785096 | PP768067 | PP768078 |
| **Gonodactyloidea** | | | | | | | | | | | | | | | |
| *Alainosquilla foresti* | PP785039 | PP785051 | PP768088 | PP796527 | PP768000 | PP768010 | PP768023 | PP785060 | PP785072 | - | PP785084 | PP768046 | PP785094 | PP768066 | PP768077 |
| *Echinosquilla guerinii* | PP785048 | PP785058 | PP768096 | PP796533 | PP768009 | PP768019 | PP768030 | PP785067 | PP785081 | PP768042 | PP785091 | - | PP785101 | PP768072 | PP768083 |
| *Gonodactylus smithii* | PP785043 | - | PP768091 | PP794581 | PP768004 | PP768015 | PP768026 | PP785064 | PP785076 | PP768037 | PP785087 | PP768050 | PP785098 | PP768069 | PP768079 |
| **Lysiosquilloidea** | | | | | | | | | | | | | | | |
| *Acanthosquilla multifasciata* | PP785037 | PP785049 | PP768086 | PP796525 | PP768098 | PP768011 | PP768022 | PP785059 | PP785070 | PP768033 | PP785082 | PP768044 | PP785093 | PP768064 | PP768075 |
| *Heterosquilla tricarinata* | PP785042 | PP785054 | PP768090 | PP796529 | PP768003 | PP768014 | PP768025 | PP785063 | PP785075 | PP768036 | PP785086 | PP768049 | PP785097 | PP768068 | - |
| *Lysiosquilla lisa* | PP785041 | PP785053 | PP768097 | - | PP768002 | PP768020 | PP768031 | PP785068 | PP785074 | PP768043 | PP785092 | PP768048 | PP785102 | PP768073 | PP768084 |
| *Tetrasquilla mccullochae* | PP785038 | PP785050 | PP768087 | PP796526 | PP768099 | PP768012 | PP768024 | PP785061 | PP785071 | PP768034 | PP785083 | PP768045 | PP785095 | PP768065 | PP768076 |
| **Squilloidea** | | | | | | | | | | | | | | | |
| *Alima maxima* | PP785045 | PP785056 | PP768093 | PP796531 | PP768006 | PP768016 | PP768027 | - | PP785078 | PP768039 | PP785089 | PP768052 | - | PP768070 | PP768080 |
| *Oratosquillina nordica* | PP785046 | - | PP768094 | PP794582 | PP768007 | PP768017 | PP768028 | PP785065 | PP785079 | PP768040 | PP785090 | PP768053 | PP785099 | - | PP768081 |
| *Squilla edentata* | PP785044 | PP785055 | PP768092 | PP796530 | PP768005 | PP768021 | PP768032 | PP785069 | PP785077 | PP768038 | PP785088 | PP768051 | PP785103 | PP768074 | PP768085 |

**Supplementary Table S3.** Matrix of stomatopod morphological characters used in this study, with character numbers and states derived from Ahyong and Harling (2000).

*The morphological data matrix will be made available upon request.*

**Supplementary Table S4.** Characters used in the morphological matrix. Names and states are derived from Ahyong and Harling (2000).

| Character number | Character name | States |
|---|---|---|
| 1 | Carapace anterolateral spines | (0) absent<br>(1) present |
| 2 | Carapace marginal carinae | (0) absent<br>(1) present |
| 3 | Carapace 'eye spots' | (0) absent<br>(1) present |
| 4 | Carapace reflected marginal carina | (0) absent<br>(1) present |
| 5 | Carapace lateral carina | (0) absent<br>(1) present |
| 6 | Carapace posterolateral angles | (0) rounded<br>(1) square<br>(2) excavate |
| 7 | Thorax lateral processes | (0) directed ventrally<br>(1) directed laterally |
| 8 | Body cross section | (0) subcylindrical<br>(1) depressed<br>(2) flattened |
| 9 | Anterolateral plate | (0) fused<br>(1) articulated |
| 10 | Abdominal submedian carinae | (0) absent<br>(1) present |
| 11 | Abdominal intermediate and lateral carinae | (0) absent<br>(1) present |
| 12 | Abdominal marginal carinae | (0) absent |

| | | |
|---|---|---|
| | | (1) present |
| 13 | Body articulation | (0) compact<br>(1) loose |
| 14 | Abdominal somite 5 lateral carina | (0) undivided<br>(1) divided |
| 15 | Abdominal somite 6 with posterolateral spine | (0) present<br>(1) absent |
| 16 | Abdominal somite 6 dorsal surface | (0) lacking upright spines<br>(1) with upright spines |
| 17 | Abdominal somite 6 articulation with telson | (0) articulated<br>(1) fused |
| 18 | Telson median surface | (0) with distinct carina<br>(1) lacking median carina<br>(2) with low broad prominence |
| 19 | Telson dorsal surface | (0) unadorned or carinate only<br>(1) tuberculate<br>(2) sculptured, irregular<br>(3) spinous |
| 20 | Postero-dorsal ornamentation above marginal armature | (0) absent<br>(1) at most irregular teeth<br>(2) regularly spaced spines<br>(3) false eave |
| 21 | Telson anterior intermediate carina | (0) absent<br>(1) present |
| 22 | Telson anterior submedian carina or boss | (0) absent<br>(1) present |
| 23 | Articulation of primary teeth of the telson | (0) all movable<br>(1) submedian teeth movable in adults<br>(2) fixed in adults |
| 24 | Telson primary teeth | (0) prominent, distinct |

| | | |
|---|---|---|
| | | (1) fused into margin forming short protrusions |
| 25 | Telson prelateral lobe | (0) absent<br>(1) present |
| 26 | Submedian denticles in adults | (0) absent<br>(1) present |
| 27 | Stridulatory carina on ventral surface of telson | (0) absent<br>(1) present |
| 28 | Intermediate denticles of the telson | (0) a maximum of 2<br>(1) absent<br>(2) 4 or more |
| 29 | Outer intermediate denticle | (0) marginal<br>(1) ventrally rotated, arising ventrally<br>(2) absent |
| 30 | Intermediate denticle shape | (0) spiniform<br>(1) rounded to triangular |
| 31 | Intermediate denticle arrangement | (0) inner on lobe<br>(1) equally and closely spaced in simple row<br>(2) with intervening lobe |
| 32 | Intervening lobe | (0) low<br>(1) produced as a denticle or short lobe<br>(2) produced as a large spine<br>(3) flattened lobe<br>(4) suppressed in adults |
| 33 | Lateral denticle | (0) absent<br>(1) present |
| 34 | Lateral denticle position | (0) marginal<br>(1) ventrally rotated |
| 35 | Lateral denticle associated lobe | (0) low |

| | | |
|---|---|---|
| | | (1) suppressed in adults<br>(2) flat, projecting lobe |
| 36 | Cornea shape | (0) globular, inclined laterally<br>(1) broadened<br>(2) bilobed, inner margin longer<br>(3) bilobed, outer margin longer |
| 37 | Ommatidia rows in cornea midband | (0) absent<br>(1) two<br>(2) three<br>(3) six |
| 38 | Corneal midband facet shape | (0) hexagonal<br>(1) rectangular<br>(2) absent |
| 39 | Cornea surface facets | (0) well defined<br>(1) poorly defined<br>(2) near absent |
| 40 | Antennal peduncle | (0) short<br>(1) long |
| 41 | Dorsal processes of antennular somite | (0) low, poorly formed<br>(1) broad, quadrate to trianguloid<br>(2) long, slender spines |
| 42 | Ventral papillae | (0) absent<br>(1) present |
| 43 | Mesial papillae | (0) absent<br>(1) present |
| 44 | Dorsal surface of antennal protopod | (0) absent or blunt<br>(1) with articulated plate<br>(2) with a fixed, laterally flattened spine |
| 45 | Antennal plate ventral surface | (0) carinate<br>(1) sulcate |

| | | |
|---|---|---|
| | | (2) fused to protopod |
| 46 | Apex of maxilla | (0) with two robust setae<br>(1) with one robust seta |
| 47 | Maxillipeds 3–4 propodi | (0) broad, with ventral ribbing<br>(1) slender |
| 48 | Maxillipeds 5 with propodal brushes | (0) present<br>(1) reduced<br>(2) vestigial |
| 49 | Raptorial claw dactylus | (0) basally uninflated<br>(1) moderately inflated<br>(2) basally inflated |
| 50 | Raptorial claw dactylus teeth size | (0) large, slender, serrated<br>(1) small, triangular |
| 51 | Raptorial claw dactylus teeth | (0) absent<br>(1) two<br>(2) three<br>(3) four<br>(4) more than four |
| 52 | Raptorial claw propodus | (0) spinous<br>(1) evenly pectinate for full length<br>(2) proximally pectinate<br>(3) fully pectinate proximally, becoming sparse distally<br>(4) smooth or sparsely pectinate |
| 53 | Raptorial claw propodal spines | (0) absent<br>(1) one<br>(2) two<br>(3) three<br>(4) four |
| 54 | Raptorial claw propodus inner margin | (0) present<br>(1) absent with setae |

| | | |
|---|---|---|
| | | (2) unadorned |
| 55 | Posterior margin of carpus of raptorial claw | (0) unarmed<br>(1) with small seta<br>(2) with distinct spine |
| 56 | Ischiomeral articulation of raptorial claw | (0) terminal<br>(1) subterminal |
| 57 | Raptorial claw ischium | (0) short<br>(1) highly reduced<br>(2) elongate<br>(3) very elongate |
| 58 | Raptorial claw ischium, lower triangular lobe | (0) absent<br>(1) present |
| 59 | Female gonopore | (0) normal<br>(1) with elaborate ornamentation |
| 60 | Pereiopodal endopod 1 | (0) slender<br>(1) elongate, ovate<br>(2) circular, ovate |
| 61 | Pereiopodal endopod 2 | (0) slender<br>(1) elongate, ovate<br>(2) circular, ovate |
| 62 | Pereiopodal endopod 3 | (0) slender<br>(1) elongate, ovate |
| 63 | Pereiopodal dactyl setation | (0) dense, soft<br>(1) sparse, robust |
| 64 | Pereiopodal endopod setation | (0) entire margin setose<br>(1) outer margin setose<br>(2) distal margin setose |
| 65 | Pleopod exopod articulated outer flap | (0) fixed<br>(1) articulated |
| 66 | Pleopod 1 endopod terminal segment setation | (0) fully setose |

| | | |
|---|---|---|
| | | (1) partially setose |
| 67 | Appendix interna | (0) slender<br>(1) stubby<br>(2) recessed |
| 68 | Pleopod endopod terminal segment | (0) large, approximately as long as broad<br>(1) reduced<br>(2) large, distinctly broader than long |
| 69 | Pleopod endopod posterior endite lateral lobe | (0) absent<br>(1) present |
| 70 | Petasma hook process apex | (0) blunt<br>(1) pointed |
| 71 | Petasma hook process | (0) elongate<br>(1) short |
| 72 | Petasma hook process | (0) broadened medially<br>(1) not broadened medially |
| 73 | Uropodal protopod primary teeth | (0) three<br>(1) two |
| 74 | Uropodal protopod inner margin | (0) smooth or crenulate<br>(1) spinous<br>(2) with minute spines |
| 75 | Uropodal endopod anterodorsal fold | (0) absent<br>(1) weak fold<br>(2) strong fold |
| 76 | Uropodal exopod segments | (0) with 2 segments<br>(1) sutured |
| 77 | Articulation of uropodal exopod segments | (0) terminal<br>(1) subterminal<br>(2) absent |

**Supplementary Table S5.** Results of Xia's saturation test in DAMBE6 (Xia, 2017) for each of the nuclear and mitochondrial genes sequenced in this study. Values are based on random 32-taxon subsamples of the complete data set. Values in bold indicate evidence of saturation.

| **Gene** | ***P*-value** | | **Iss** | **Iss.c** | |
|---|---|---|---|---|---|
| | **asymmetrical tree** | **symmetrical tree** | | **asymmetrical tree** | **symmetrical tree** |
| Nuclear *18S* | <0.00005 | <0.00005 | 0.106 | 0.518 | 0.780 |
| Nuclear *28S* | <0.00005 | <0.00005 | 0.716 | **0.542** | 0.804 |
| Nuclear *H3* | <0.00005 | <0.00005 | 0.153 | 0.418 | 0.694 |
| $1^{st}$ codon sites of mitochondrial protein-coding genes | <0.00005 | <0.00005 | 0.497 | 0.552 | 0.808 |
| $2^{nd}$ codon sites of mitochondrial protein-coding genes | <0.00005 | <0.00005 | 0.431 | 0.552 | 0.808 |
| $3^{rd}$ codon sites of mitochondrial protein-coding genes | <0.00005 | <0.00005 | 0.956 | **0.552** | **0.808** |
| Mitochondrial *12S* | <0.00005 | 0.7546 | 0.726 | **0.417** | 0.734 |
| Mitochondrial *16S* | <0.00005 | <0.00005 | 0.877 | **0.481** | **0.769** |
| Mitochondrial tRNA genes | <0.00005 | <0.00005 | 0.392 | 0.489 | 0.773 |

**Supplementary Table S6.** Nine-subset partitioning scheme implemented in all stomatopod phylogenetic analyses.

| **Gene** | **Nucleotides** |
|---|---|
| Nuclear *18S* | 1825 |
| Nuclear *28S* | 2898 |
| Nuclear *H3* | 411 |
| Mitochondrial *12S* | 824 |
| Mitochondrial tRNAs | 1470 |
| +ve strand 1$^{st}$ codon sites of mitochondrial protein-coding genes | 2241 |
| +ve strand 2$^{nd}$ codon sites of mitochondrial protein-coding genes | 1344 |
| -ve strand 1$^{st}$ codon sites of mitochondrial protein-coding genes | 2241 |
| -ve strand 2$^{nd}$ codon sites of mitochondrial protein-coding genes | 1344 |

**Supplementary Table S7.** List of fossils used to calibrate nodes in the molecular dating analyses of Stomatopoda. All calibration priors include a 97.5% soft maximum based on the age of crown stomatopods as suggested by the appearance of *Daidal acanthocercus* Jenner, Hof & Schram, 1998 (Stomatopoda, Archaeostomatopodea) in the Carboniferous (313 million years ago) (Schram, 2007; Van Der Wal et al., 2017).

| **Fossil taxon** | **Node (Genus)** | **Exponential prior (millions of years)** | | **Reference** |
|---|---|---|---|---|
| | | **Offset** | **Mean** | |
| ***Pseudosquilla berica* De Angeli & Messina, 1996**<br><br>(Stomatopoda, Gonodactyloidea) | *Pseudosquilla*<br>+<br>*Raoulserenea* | 33.90 | 75.65 | De Angeli and Messina (1996) |
| ***Odontodactylus italicus* Beschin, De Angeli, Checchi & Zarantonello, 2012**<br><br>(Stomatopoda, Gonodactyloidea) | *Gonodactylus*, *Gonodactylaceus*, *Chorisquilla*, and *Echinosquilla*<br>+<br>*Odontodactylus* | 41.20 | 73.70 | Beschin et al. (2012) |
| ***Bathysquilla wetherelli* (Woodward, 1879)**<br><br>(Stomatopoda, Bathysquilloidea) | *Bathysquilla*<br>+<br>*Indosquilla* | 22.98 | 78.60 | Schram et al. (2013) |
| ***Squilla taulinanus* Ahyong, Charbonnier & Garassino, 2013**<br><br>(Stomatopoda, Squilloidea) | *Squilla*<br>+<br>*Harpiosquilla* | 15.92 | 80.50 | Ahyong et al. (2013) |

| | | | | |
|---|---|---|---|---|
| ***Ursquilla yehoachi* (Remy & Avnimelech, 1955)**<br><br>(Stomatopoda, Squilloidea) | *Alima*, *Harpiosquilla*, *Kempella*, *Oratosquilla*, *Oratosquillina*, *Quollastria*, *Squilla*, and *Squilloides*<br>+<br>*Alainosquilla foresti* | 63.40 | 79.01 | Haug et al. (2013) |

**References**


Ahyong, S.T., Charbonnier, S., Garassino, A., 2013. Squilla taulinanus n. sp (Crustacea, Stomatopoda, Squillidae) from the Burdigalian (Miocene) of Taulignan, south-eastern France. Bol. Soc. Geol. Mex. 65, 213–217.

Ahyong, S.T., Harling, C., 2000. The phylogeny of the stomatopod Crustacea. Aust. J. Zool. 48, 607–642.

Beschin, C., De Angeli, A., Checchi, A. & Zarantonello, G., 2012. Crostacei del giacimento Eocenico di Grola presso spagnado di Cornedo Vicentino (Vicenza, Italia settentrionale) (Decapoda, Stomatopoda, Isopoda). Museo di Archeologia e Scienze Naturali “G. Zannato”, Montecchio Maggiore, Italy.

De Angeli, A., Messina, V., 1996. Pseudosquilla berica nuova specie di Stomatopoda del Terziario Veneto. Studi e ricerche, Associazione Amici del Museo Civico 6, 5–10.

Haug, C., Kutschera, V., Ahyong, S. T., Vega, F. J., Maas, A., Waloszek, D., Haug, J. T., 2013. Re-evaluation of the Mesozoic mantis shrimp *Ursquilla yehoachi* based on new material and the virtual peel technique. Palaeontol. Electron., 16, 5T1–14.

Schram, F.R., 2007. Paleozoic proto-mantis shrimp revisited. J. Paleontol. 81, 895–916.

Schram, F.R., Klein, J.C.v.V., Charmantier-Daures, M., 2013. Treatise on Zoology - Anatomy, Taxonomy, Biology. The Crustacea, Volume 4 Part A. Brill, Leiden, The Netherlands.

Van Der Wal, C., Ahyong, S.T., Ho, S.Y.W., Lo, N., 2017. The evolutionary history of Stomatopoda (Crustacea: Malacostraca) inferred from molecular data. PeerJ 5, e3844.

Xia, X., 2017. DAMBE6: New tools for microbial genomics, phylogenetics, and molecular evolution. J. Hered. 108, 431–437.
